# Prognostic implications and biological roles of EMT-related lncRNAs in lung squamous cell carcinoma: an in-depth analysis utilizing a novel prognostic signature and classification system

**DOI:** 10.1101/2023.10.03.560645

**Authors:** Jinming Zhang, Baihong Zheng, Xiuying Zhang, Ying Liu, Ying Guo, Jia Zhao, Jiamei Liu, Hui Xue

## Abstract

**Background:** Lung squamous cell carcinoma (LUSC) represents a major subtype of non-small cell lung cancer (NSCLC), a leading contributor to cancer-related mortality. Epithelial-mesenchymal transition (EMT)-associated genes have been implicated in poor survival and metastatic gene expression in LUSC. Long non-coding RNAs (lncRNAs) are known to facilitate tumor progression and metastasis via EMT regulation. However, the prognostic significance and biological functions of EMT-associated lncRNAs in LUSC remain to be elucidated.

**Methods:** In this study, we aimed to develop an EMT-related lncRNA prognostic signature (EMT-LPS) utilizing RNA transcription data from LUSC patients in The Cancer Genome Atlas (TCGA) database, along with corresponding clinical characteristics. Kaplan-Meier analysis, receiver operating characteristic (ROC) curves, and Cox regression were employed to validate and assess the model. Furthermore, we confirmed the independent prognostic value of key genes in EMT-LPS using Gene Expression Profiling Interactive Analysis (GEPIA). Additionally, we proposed a novel LUSC classification system based on EMT-related lncRNA expression patterns, evaluating the prognostic profile, tumor microenvironment, and immunotherapy sensitivity of each subtype.

**Results:** A prognostic signature comprising twelve genes was constructed, and patients were stratified into high and low-risk groups according to their risk scores. Cox regression analysis revealed that the risk score served as an independent prognostic factor. A nomogram was generated to predict LUSC patient survival rates. Distinct subtypes exhibited varying tumor purity, immunogenicity, and immunotherapy drug sensitivity.

**Conclusions:** Our findings underscore the relevance of EMT-related lncRNAs in LUSC and their potential utility in guiding immunotherapy strategies. The EMT-LPS and novel LUSC typing scheme provide a new perspective for understanding the biological functions and prognostic role of EMT-related lncRNAs in LUSC.

## 1 Introduction

Lung squamous cell carcinoma (LUSC) represents the second most prevalent subtype of non-small cell lung cancer (NSCLC) (1), a leading cause of cancer-related mortality worldwide, accounting for 85% of all lung cancer cases. This high mortality rate is attributed to late-stage diagnosis, rapid metastasis, and recurrence (2). Despite comprehensive treatment approaches involving surgery, radiotherapy, and chemotherapy, the 5-year survival rate for LUSC remains a dismal 5% (3), with limited therapeutic options available. Recent advancements in immunotherapy have expanded the treatment repertoire for LUSC (4); however, its clinical application remains restricted, and the criteria for its use warrant further refinement (5). Consequently, the identification of novel biomarkers or therapeutic targets for predicting LUSC progression, prognosis, and treatment response, particularly to immunotherapy, is of paramount importance.

Epithelial-mesenchymal transition (EMT) is a cellular process that occurs widely in physiological or pathological processes, playing a crucial role in tumor progression (6). EMT represents a reversible dedifferentiation process, whereby activated tumor cells lose epithelial-like properties, such as intercellular adhesion and polarity, resulting in enhanced migration and invasion. Concurrently, tumor cells acquire mesenchymal stem cell characteristics, facilitating therapeutic resistance and ultimately leading to tumor recurrence and metastasis (7). Notably, EMT-related CDR1 has been implicated in poor survival and the expression of metastatic gene signatures in LUSC (8), suggesting that EMT-associated genes may serve as promising therapeutic targets. Although long non-coding RNAs (lncRNAs) have been demonstrated to regulate EMT and promote tumor progression and metastasis (9), the relationship between EMT-related lncRNA expression profiles, LUSC clinicopathological features, and patient prognosis remains unexplored.

In this study, we utilized transcriptomic data from LUSC patients in The Cancer Genome Atlas Program (TCGA) database to construct a robust EMT-related lncRNA signature for predicting patient prognosis. Additionally, we assessed the prognostic significance of potential EMT-related lncRNA subtypes in LUSC patients and examined the associations between these subtypes, the tumor microenvironment, and immunotherapy response.

## 2 Materials and methods

### 2.1 Data collection

For this study, we obtained RNA sequencing files of LUSC patients from the TCGA database (https://portal.gdc.cancer.gov), which were standardized as Fragments Per Kilobase of exon model per Million mapped fragments (FPKM). Corresponding clinicopathological information was also collected. Patients with missing overall survival (OS) data were excluded, resulting in a total of 495 LUSC patients for downstream analysis. To identify EMT-associated genes, we utilized the Molecular Signatures Database (MsigDB) v7.4 (https://www.gsea-msigdb.org/gsea/msigdb) and selected the marker gene set named HALLMARKEPITHELIALMESENCHYMAL_TRANSITION. We collected 200 EMT-associated genes, which are listed in **S1 Table**.

### 2.2 Selection of EMT-related lncRNAs

In accordance with the human genome annotation files available on the GENCODE website (https://www.gencodegenes.org/human), we extracted all lncRNA expression data from the aforementioned RNA sequencing data. Subsequently, we employed Pearson correlation analysis to identify EMT-related lncRNAs, utilizing the filtering criteria of |CorPearson|>0.3 and *p-*values <0.01.

### 2.3 Development of a lncRNA-based prognostic signature

Firstly, the expression data of EMT-related lncRNAs were integrated with patient survival data and subjected to univariate Cox regression analysis to identify lncRNAs that are significantly associated with prognosis.

Next, we randomly divided a cohort of 495 LUSC patients into two groups: a TRAIN set (n=297) and a TEST set (n=198). The expression data of prognostic lncRNAs were analyzed using Least Absolute Shrinkage and Selection Operator (LASSO) regression with the R package “glmnet”. The penalty parameter was optimized using cross-validation.

Ultimately, we developed an EMT-related lncRNA prognostic signature (EMT-LPS) for LUSC patients, which consists of EMT-associated lncRNAs. The risk score of each patient can be calculated using the following equation:

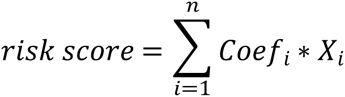

where X_i_ represents the FPKM value of each prognostic lncRNA and Coef_i_ represents the coefficient of each prognostic lncRNA.

### 2.4 Analysis of the prognostic value of lncRNA prognostic signature

Patients were stratified into high-risk and low-risk groups based on their risk scores. Subsequently, receiver operating characteristic (ROC) curves, survival curves, and prognostic signature lncRNA expression heatmaps were generated for both the TRAIN and TEST sets to assess the predictive accuracy of the EMT-LPS. Univariate Cox regression and multivariate Cox regression analyses were employed to ascertain if EMT-LPS and clinical characteristics could serve as independent prognostic predictors for LUSC patients. To further investigate the prognostic value of EMT-LPS across various clinical variable strata, patients were categorized according to age, gender, stage, and TNM classification. Survival curves based on risk scores were constructed for each group separately. A *p-* value of <0.05 was considered statistically significant in the aforementioned analyses.

### 2.5 Construction and validation of nomogram

The R package “rms” was utilized to develop a nomogram incorporating patients’ stage and risk scores. The ROC curve was then employed to evaluate the nomogram’s performance.

### 2.6 Gene set enrichment analysis (GSEA)

To examine differences in underlying biological processes and pathways between high-risk and low-risk subgroups, GSEA was conducted separately for both subgroups. A *p-*value of <0.05 and a False Discovery Rate (FDR) *q-*value of <0.05 were deemed statistically significant.

### 2.7 Consensus clustering

The R package “ConsensusClusterPlus” was employed to perform consensus clustering on LUSC samples to uncover potential molecular subtypes. The number of clusters (k) was set from 2 to 9, and the optimal k value was chosen based on the empirical cumulative distribution function (CDF) plot and the delta area plot curve to obtain a suitable classification. Subsequently, mutations of the top 20 genes with the highest mutation rate in LUSC were analyzed for each cluster and visualized using waterfall plots generated by the R package “maftools”.

### 2.8 Evaluation of tumor microenvironment

To assess the tumor microenvironment in LUSC, the CIBERSORT algorithm was employed to estimate the infiltration levels of 22 immune cell types based on whole-gene expression levels, running the algorithm for 1000 permutations using the LM22 signature. Samples with output *p-* values of <0.05 were selected for downstream analyses. The stromal score, immune score, ESTIMATE score, and tumor purity were then calculated for each sample using the R package “ESTIMATE”.

### 2.9 Analysis of immunotherapy response

Leveraging immunotherapy information such as Immunophenoscore (IPS) for LUSC patients provided by the Cancer Immunome Atlas (TCIA) database, the efficacy of various immunotherapies among LUSC patients with different clusters was analyzed. Immunotherapy sensitivity data were visualized using violin plots generated with the R package “ggpubr”. A *p-*value of <0.05 was considered indicative of a statistically significant difference.

### 2.10 Key genes validation

Gene Expression Profiling Interactive Analysis (GEPIA, http://gepia.cancer-pku.cn/) is a novel web-based tool that incorporates sequencing expression data from 9,736 tumor samples and 8,587 normal samples across 33 cancer types. In this study, we employed GEPIA to investigate the independent impact of key genes that are positively associated with tumor progression in the prognostic model on the prognosis of patients with LUSC (screening criteria: coefficient > 1). We utilized default parameters, with the group cutoff set at “median” and hazard ratio (HR) and 95% confidence interval (CI) set at “yes”.

### 2.11 Statistical analyses

The statistical analysis for this study was conducted utilizing the R programming language (version 4.0.3) and SPSS Statistics 25 software. Continuous variables were assessed using the Wilcoxon test, while categorical variables were examined through Fisher’s exact test or the chi-square test, as appropriate. Survival differences were evaluated using Kaplan-Meier curves and log-rank tests. A *p-* value of less than 0.05 was deemed to indicate statistical significance.

### 2.12 Ethical review

In this study, all the research data were obtained from publicly accessible databases, namely TCGA, GEPIA, and MsigDB. These databases provide de-identified, pre-existing, publicly available data, and therefore, the use of such data does not constitute human subjects research. Consequently, the study did not require review or approval from an ethics committee. The data extraction and analysis were performed in compliance with the terms and conditions of the respective databases.

## 3 Results

### 3.1 Identification of EMT-related lncRNAs

A total of 14,086 lncRNA genes were extracted from RNA-seq files obtained from the TCGA database, utilizing the human genome annotation file from the GENCODE website. Concurrently, 200 EMT-related genes were gathered from the MSigDB database for subsequent analysis. A correlation analysis was conducted between the lncRNA expression of the acquired samples and their EMT gene expression. lncRNAs associated with one or more EMT genes were identified as EMT-related lncRNAs, based on the screening criteria of |CorPearson|>0.3 and *p-*values <0.01. The final set of 2,036 EMT-associated lncRNAs is provided in **S2 Table**.

### 3.2 Construction of EMT-LPS

Univariate Cox regression analysis between EMT-related lncRNA expression data and patients’ clinical data yielded 107 lncRNAs significantly associated with prognosis. The 495 patients included in the study were randomly divided into TRAIN and TEST sets. Lasso regression analysis was performed in the TRAIN set to construct an EMT-LPS comprising 12 lncRNAs (**Fig 1**). The risk score of LUSC patients could be calculated using the following penalty function formula:

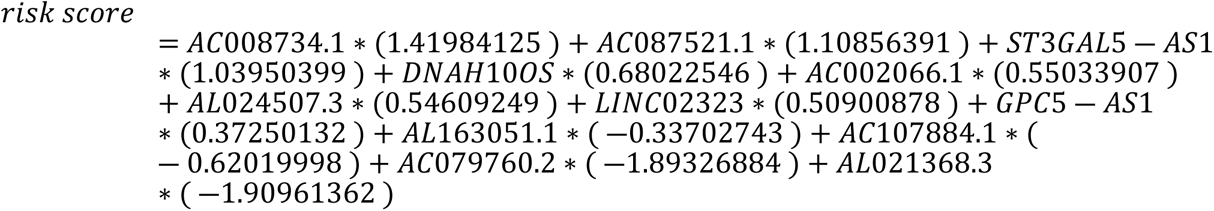

**Fig 1.**
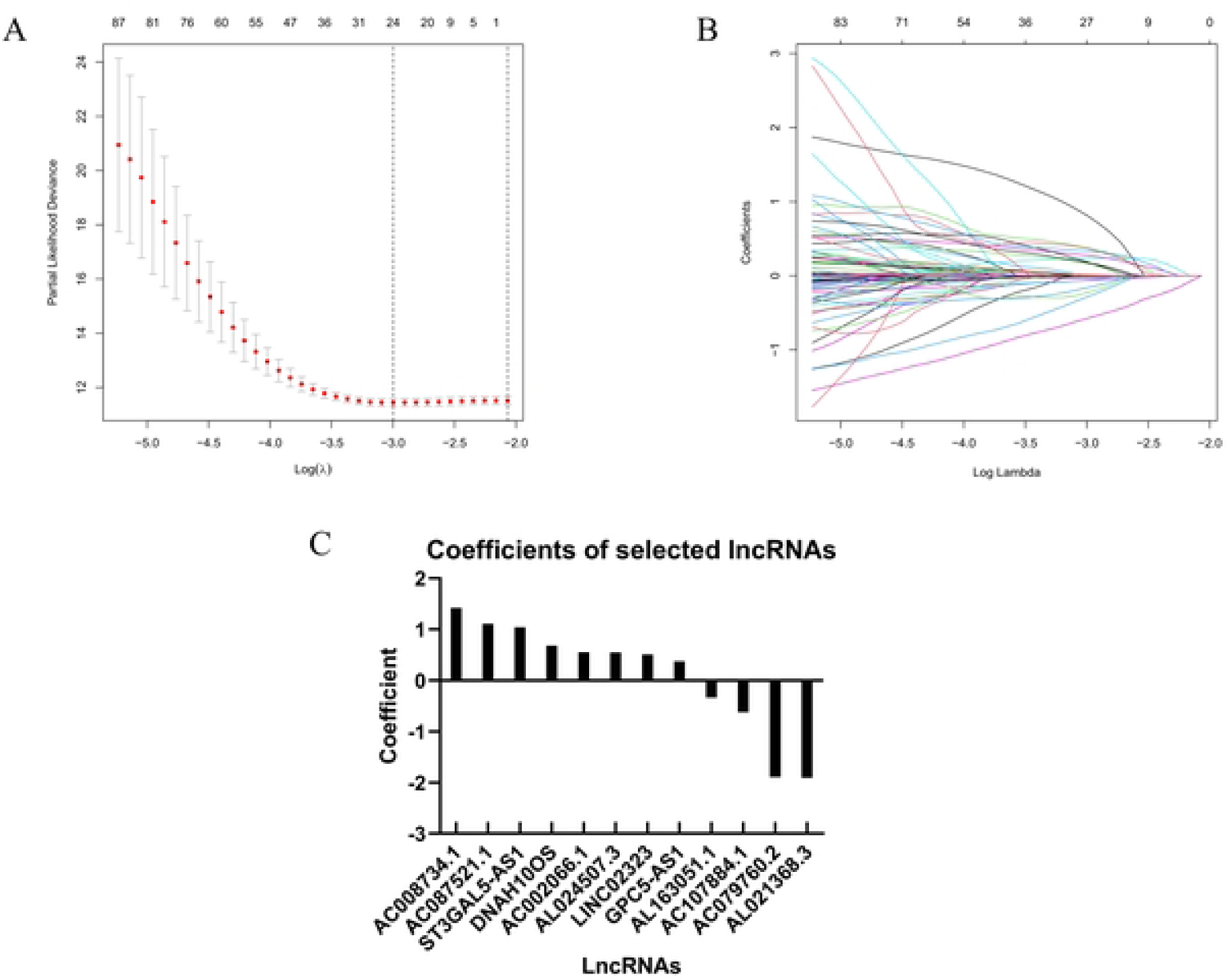
The optimal model obtained through LASSO regression (A, B) and the corresponding coefficients of the selected lncRNAs (C) are presented herein.

### 3.3 Evaluation of the prognostic predictive value of EMT-LPS in LUSC

Upon determining the risk score for each sample based on the derived EMT-LPS, patients were stratified into high-risk and low-risk groups using the median risk score. Kaplan-Meier survival curves were generated for both the TRAIN and TEST sets to evaluate potential differences in OS between the high- and low-risk groups (**Fig 2A, B**). The results demonstrated a significant disparity in OS between patients in the high-risk group compared to those in the low-risk group within both datasets, with patients in the low-risk group exhibiting longer survival times. ROC curves were also constructed with 1, 2, and 3 years as the defined time points. The area under the curve (AUC) values for the TRAIN set were 0.722, 0.752, and 0.775, respectively (**Fig 2C**), while the AUC values for the TEST set were 0.647, 0.710, and 0.682 in the same order (**Fig 2D**). These findings indicate the strong prognostic prediction accuracy of the EMT-LPS, particularly in patients with extended prognostic survival times.

**Fig 2.**
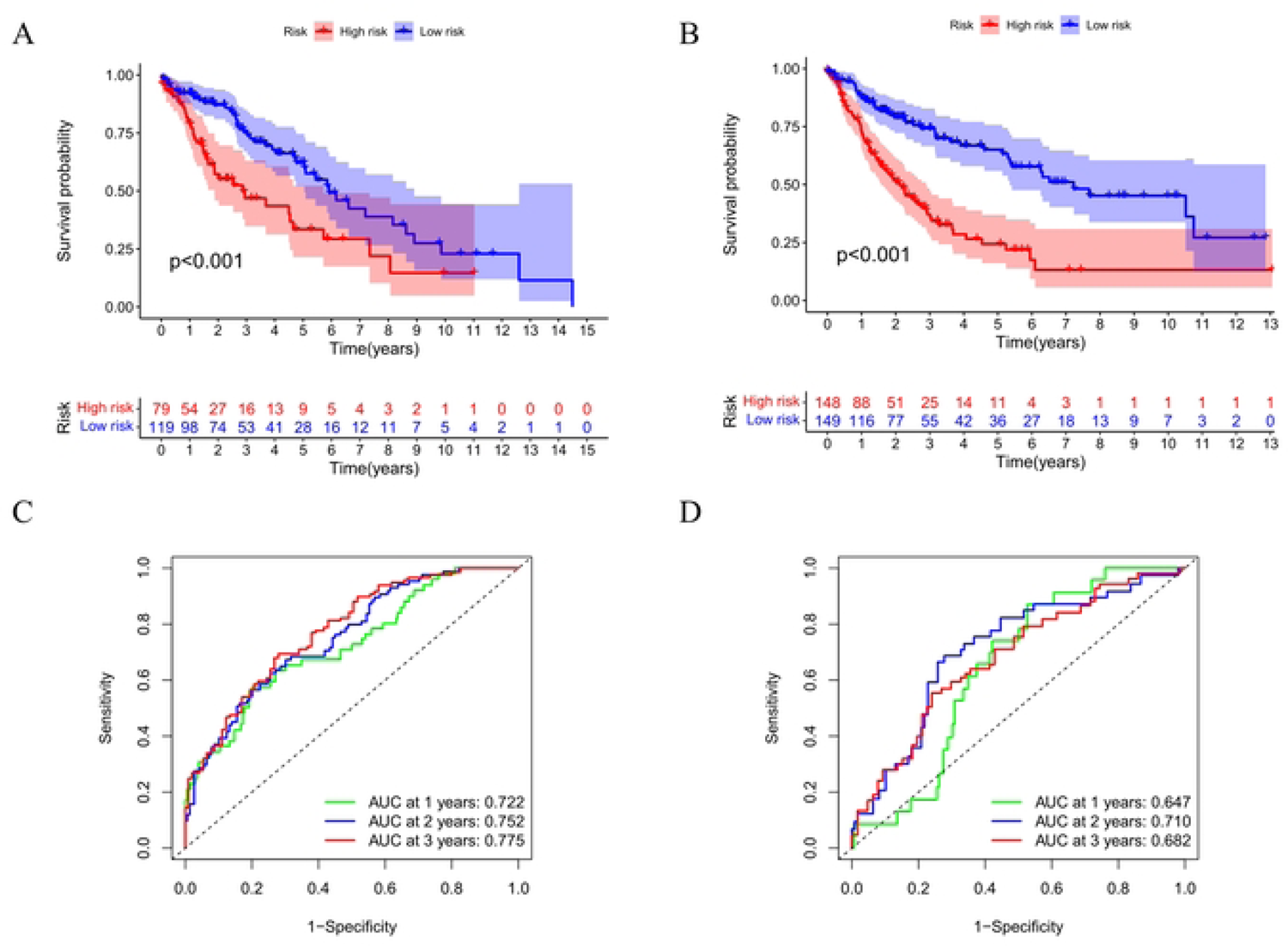
The prognostic value of EMT-LPS in LUSC. Kaplan-Meier survival curves revealed that the high-risk group exhibited poorer overall survival compared to the low-risk group in both the TRAIN (A) and TEST (B) sets. Receiver operating characteristic curves were generated to evaluate the ability of EMT-LPS to predict 1/2/3-year survival in the TRAIN (C) and TEST (D) sets.

The trends in risk scores, distribution of survival status, and expression of the 12 lncRNAs encompassed within the EMT-LPS are depicted in **Figs 3A, B**. The data revealed that AC008734.1, AC087521.1, ST3GAL5-AS1, DNAH10OS, AC002066.1, AL024507.3, LINC02323, and GPC5-AS1 were predominantly associated with elevated risk scores and increased likelihood of mortality, whereas high expression of genes such as AL163051.1, AC107884.1, AC079760.2, and AL021368.3 were frequently observed in samples with lower risk scores.

**Fig 3.**
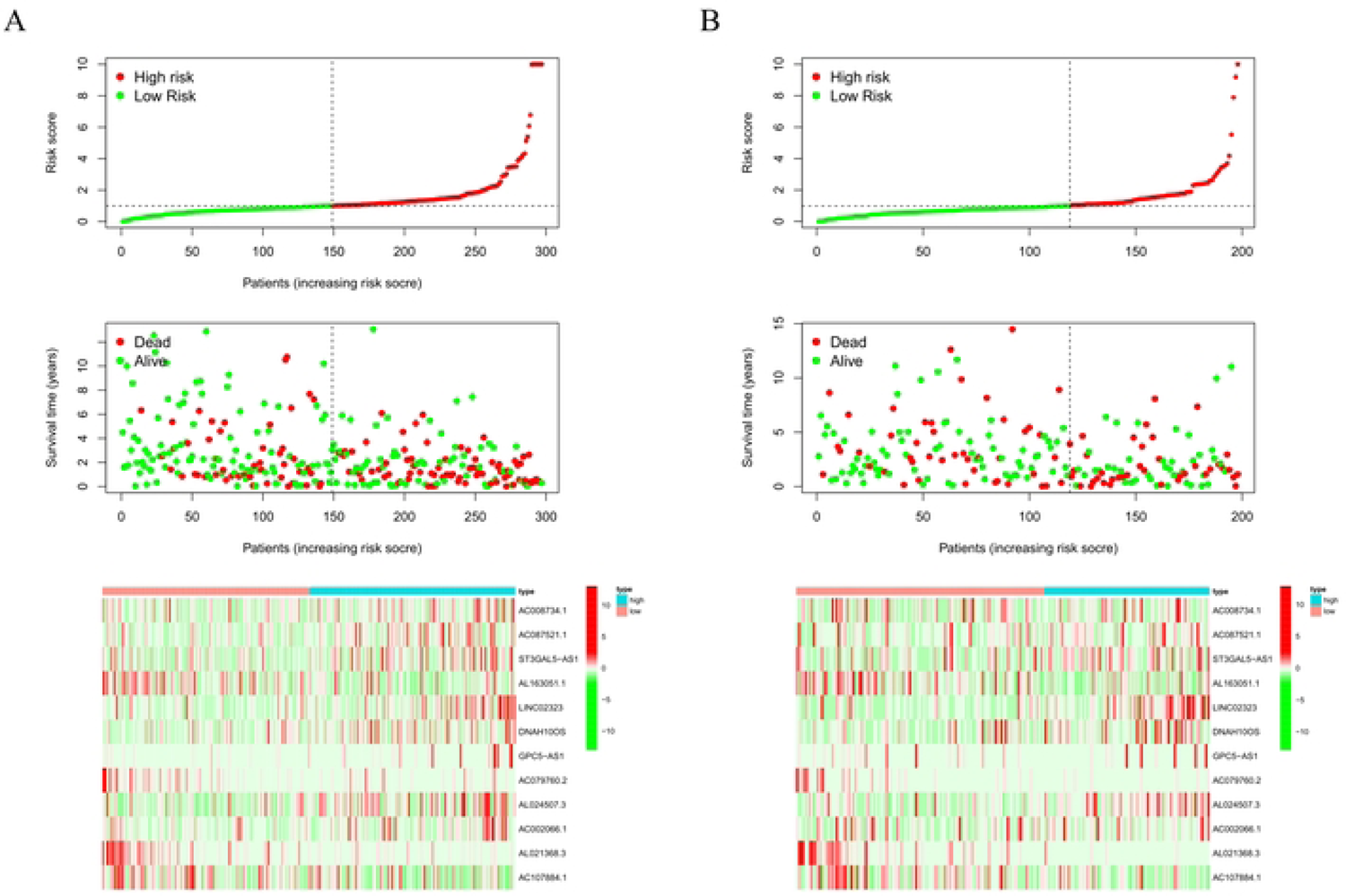
The trends of risk scores, the distribution of survival status, and the expression of 12 lncRNAs included in the EMT-LPS in both the TRAIN (A) and TEST (B) sets.

Subsequent analysis of prognostic correlates was conducted on the entire cohort of 495 LUSC patients. Univariate Cox regression analysis revealed that age, pathological stage, and risk score were strongly associated with patient prognosis (**Fig 4A**). Multivariate Cox regression analysis demonstrated that both pathological stage and risk score were independent predictors of prognosis in LUSC patients (**Fig 4B**).

**Fig 4.**
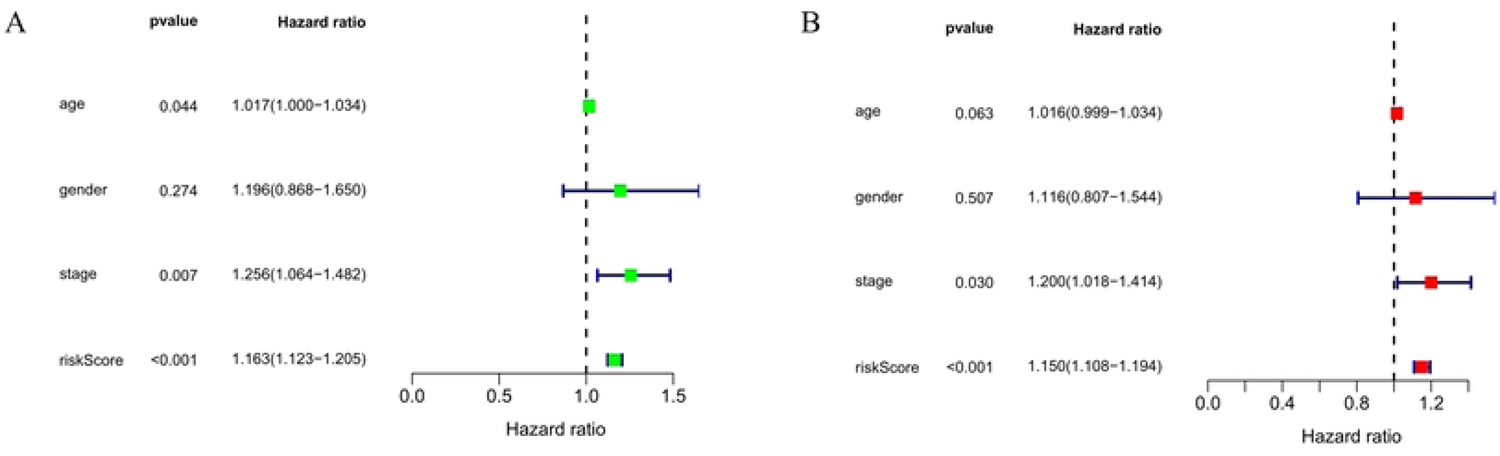
Utilizing univariate Cox regression (A) and multivariate Cox regression (B), the independent predictive ability of risk scores and clinical features for prognosis were analyzed.

The applicability of EMT-LPS across various clinical variable strata was further investigated (**Fig 5**). Kaplan-Meier survival curves indicated that, with the exception of stage M1 LUSC patients with distant tumor metastases (comprising only 7 cases, resulting in excessive statistical error due to the small sample size), significant survival differences were observed between the high-risk and low-risk groups across most clinical variables, including age, gender, pathological grade, and TNM stage. Patients in the high-risk group consistently exhibited shorter survival times, thereby confirming the robustness of the prognostic function of EMT-LPS.

**Fig 5.**
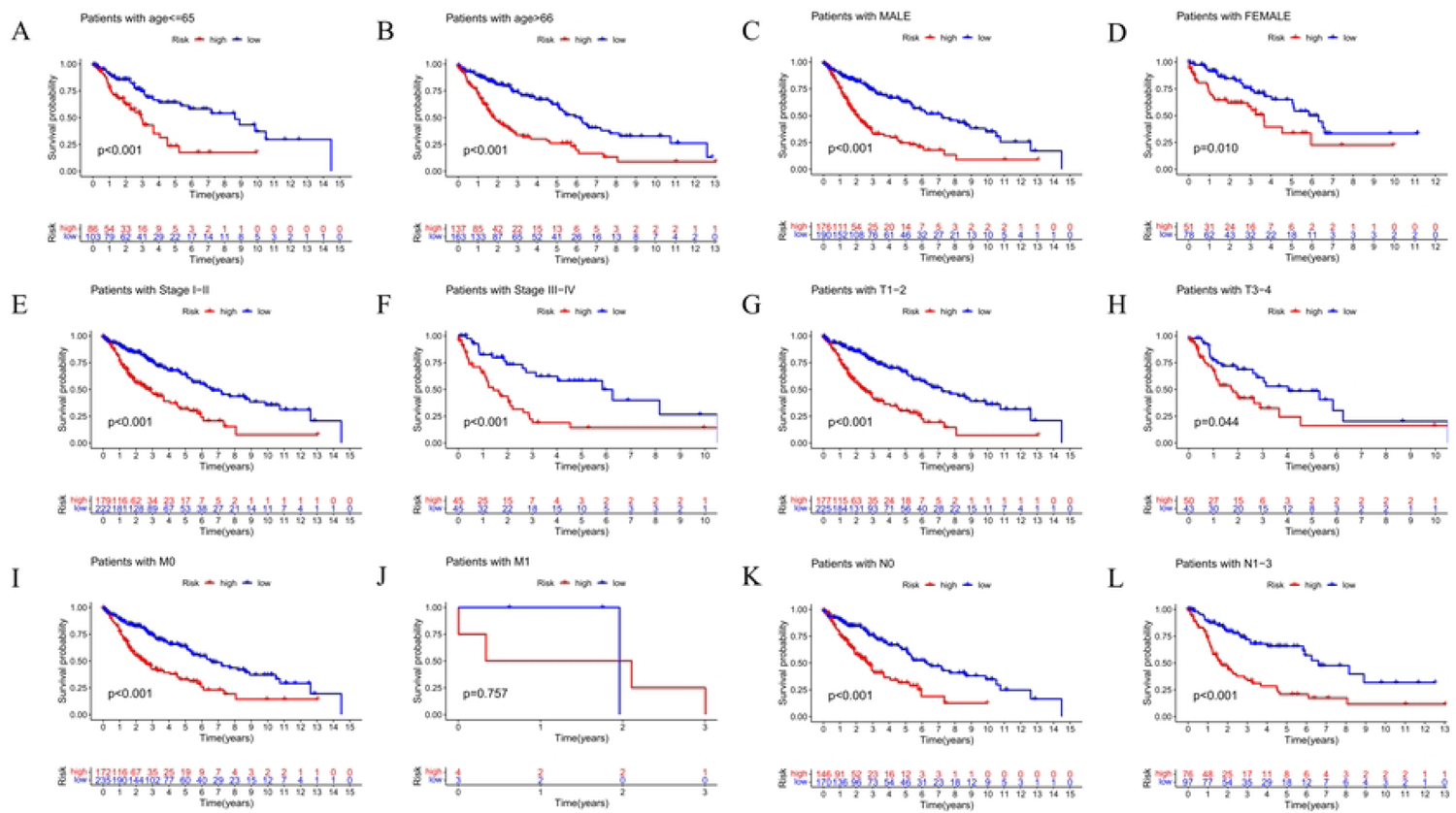
**Survival analysis was conducted for the high-risk and low-risk groups** stratified by various clinical features, including age (A, B), gender (C, D), stage (E, F), tumor (G, H), lymph node (I, J), and metastasis (K, L).

### 3.4 Development and validation of the nomogram

Utilizing the data acquired from the entire cohort, a nomogram was constructed to prognosticate the survival duration of patients based on their tumor stage and EMT-LPS risk score (**Fig 6A**). The performance of the developed nomogram was assessed using ROC curves, and the analysis revealed that AUC values for the 1-year, 2-year, and 3-year ROC curves were 0.650, 0.692, and 0.704, respectively **(Fig 6B**). These results indicate that the constructed nomogram demonstrates a relatively precise prognostic ability for OS in the field of LUSC.

**Fig 6.**
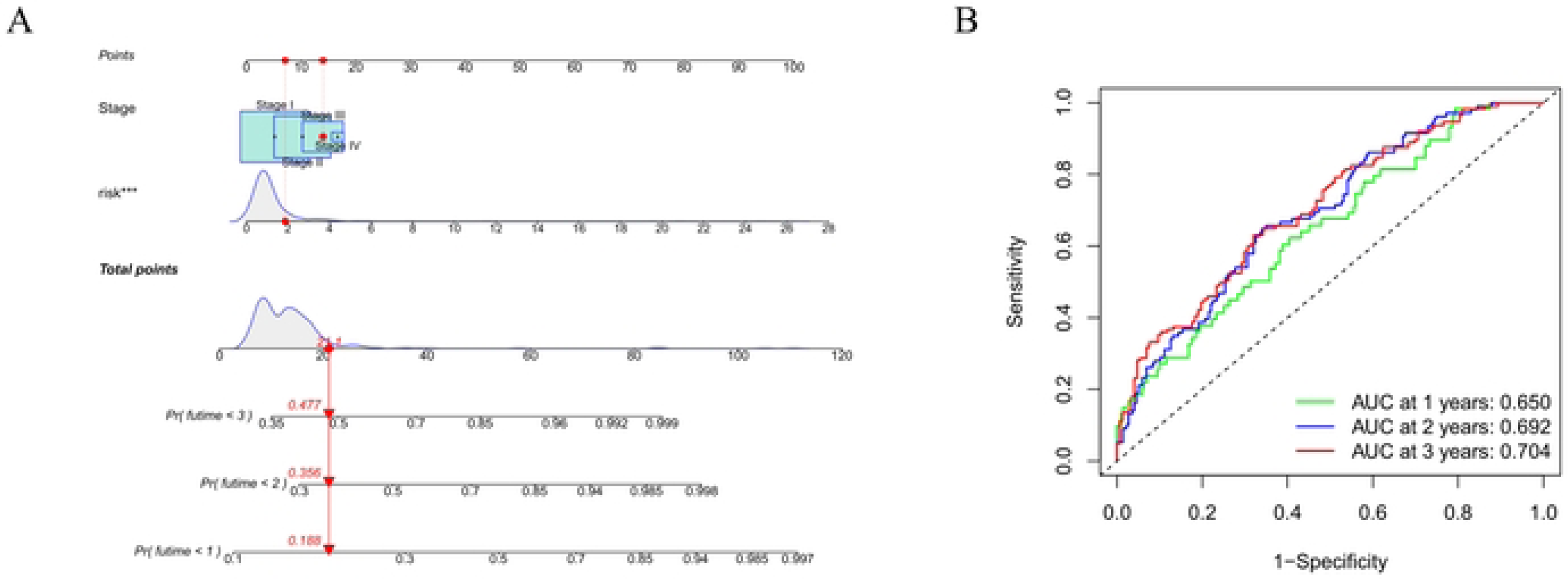
**A nomogram was developed to predict the overall survival of patients based on their stage and risk score of EMT-LPS (A). Receiver operating characteristic curves were generated to evaluate the predictive performance of the nomogram for 1/2/3-year survival (B).**

### 3.5 Enrichment pathway analysis using GSEA

We employed GSEA to investigate the enriched metabolic pathways in high- and low-risk groups. All signaling pathways with an FDR *q-*value < 0.05 are presented in **Table 1**, along with their related information. Furthermore, we selected several signaling pathways exhibiting the most significant differences between high- and low-risk groups and depicted their details in **Fig 7**. In the high-risk group, the predominantly enriched pathways encompassed extracellular matrix (ECM) receptor interaction, focal adhesion, leukocyte transendothelial migration, cytokine-cytokine receptor interaction, nucleotide-binding oligomerization domain (NOD)-like receptor signaling pathway, and Janus kinase-signal transducer and activator of transcription (JAK-STAT) signaling pathway, among others (**Fig 7A**). Conversely, the low-risk group primarily exhibited enrichment in pathways related to Alzheimer’s disease, oxidative phosphorylation, and Parkinson’s disease (**Fig 7B**). These pathways may contribute to a deeper understanding of the biological alterations occurring in patients with lung squamous cell carcinoma.

**Fig 7.**
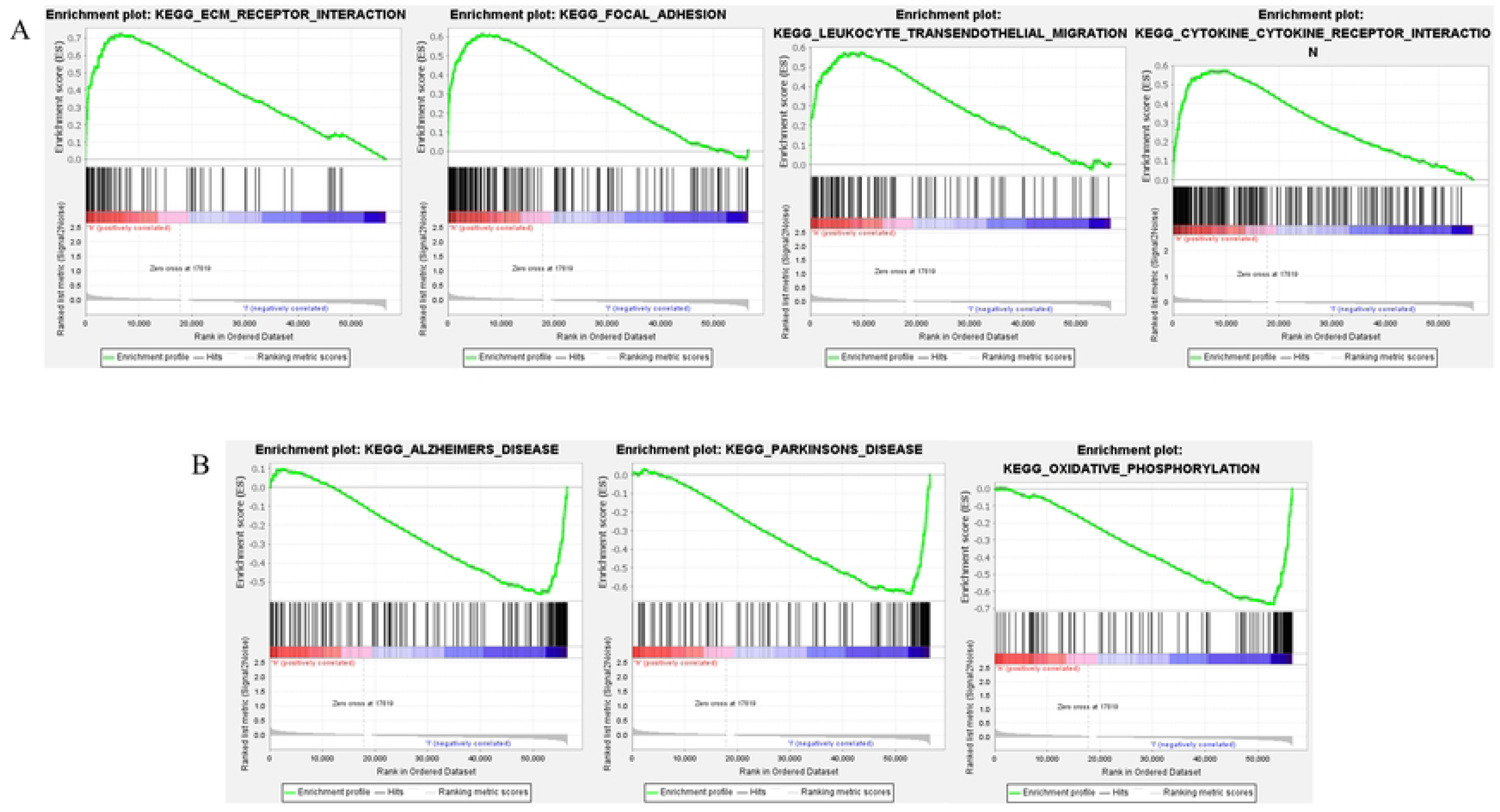
The results of the GSEA enrichment pathway analysis, highlighting the selected gene pathways that were enriched in the high-risk (A) and low-risk (B) groups.

**Table 1.** Gene set enrichment analysis results.

| Gene Set | Size | Upregulated in group | Normalized enrichment score | FDR $q$-value |
| --- | --- | --- | --- | --- |
| ECM RECEPTOR INTERACTION | 84 | High-risk group | 2.15 | 0.009 |
| FOCAL ADHESION | 199 | High-risk group | 2.17 | 0.015 |
| LEUKOCYTE TRANSENDOTHELIAL MIGRATION | 116 | High-risk group | 2.03 | 0.019 |
| CYTOKINE-CYTOKINE RECEPTOR INTERACTION | 264 | High-risk group | 2.04 | 0.020 |
| NOD-LIKE RECEPTOR SIGNALING PATHWAY | 62 | High-risk group | 2.06 | 0.022 |
| JAK-STAT SIGNALING PATHWAY | 155 | High-risk group | 1.99 | 0.022 |
| HYPERTROPHIC CARDIOMYOPATHY HCM | 83 | High-risk group | 1.96 | 0.022 |
| DORSO VENTRAL AXIS FORMATION | 24 | High-risk group | 1.96 | 0.024 |
| COMPLEMENT AND COAGULATION CASCADES | 69 | High-risk group | 2.00 | 0.025 |
| ARRHYTHMOGENIC RIGHT VENTRICULAR CARDIOMYOPATHY ARVC | 74 | High-risk group | 1.96 | 0.026 |
| DILATED CARDIOMYOPATHY | 90 | High-risk group | 1.88 | 0.041 |
| ALZHEIMERS DISEASE | 165 | Low-risk group | -2.03 | 0.030 |
| OXIDATIVE PHOSPHORYLATION | 131 | Low-risk group | -2.04 | 0.043 |
| PARKINSONS DISEASE | 128 | Low-risk group | -1.97 | 0.046 |

### 3.6 Classification of LUSC subtypes based on EMT-Associated lncRNAs

We investigated the potential molecular subtypes of LUSC using the Consensus Clustering algorithm, which was applied to the expression data of EMT-related lncRNAs significantly associated with prognosis across all cases. The resulting classification is presented in Fig 8. The cumulative distribution function (CDF) plot (**Fig 8A**) indicated that the distribution reached an approximate maximum when k=7. Additionally, the delta area plot (**Fig 8B**) demonstrated a minor increase in the distribution when k=8. Based on the rule (10) of the R package “ConsensusClusterPlus”, we determined that the optimal clustering result for LUSC was achieved at k=7, with the consensus matrix plot (**Fig 8C**) displaying the “cleanest” cluster partition at this point. Nonetheless, considering the inadequate sample size in three of the clusters (three cases in Cluster 5, one case in Cluster 6, and two cases in Cluster 7), along with the tracking plot (**S1 Fig**) indicating weak class membership for these six cases, it was deemed necessary to enhance generalizability and scalability for subsequent analyses. As a result, we incorporated 489 patients from Clusters 1, 2, 3, and 4 into the downstream study, while excluding the six patients from Clusters 5, 6, and 7.

**Fig 8.**
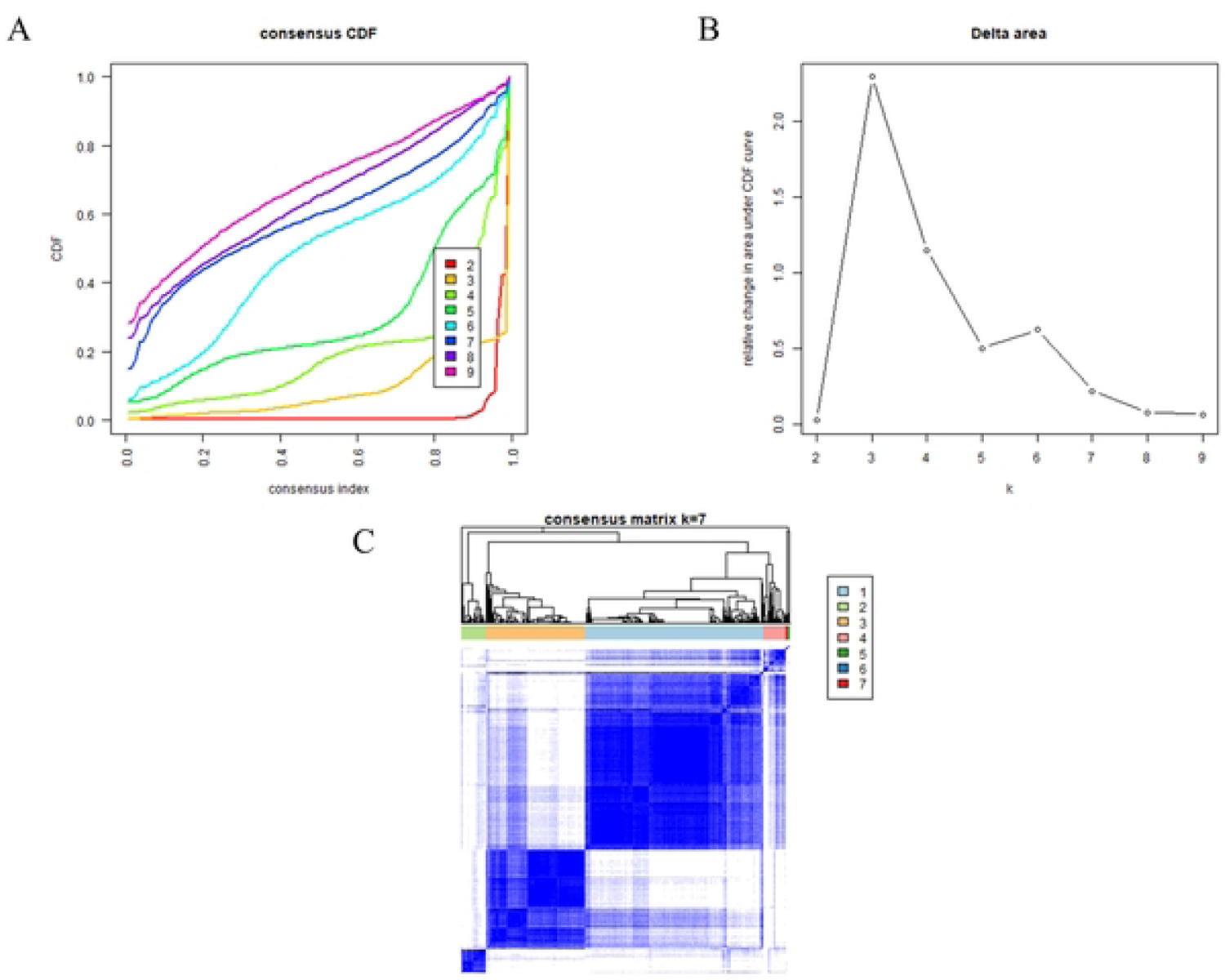
Consensus clustering algorithm analysis of potential EMT-related LUSC subtypes. Empirical cumulative distribution function plot (A). Delta area plot curve (B). Consensus matrix plot at k=7 (C).

### 3.7 Clinical features and genetic mutation characteristics in different subtypes

Kaplan-Meier survival analysis was performed for the four subtypes, revealing distinct OS outcomes among them (**Fig 9**). Cluster 3 was generally associated with longer survival times and higher survival rates, whereas Cluster 4 exhibited poorer OS compared to the other subtypes. Clusters 1 and 2 demonstrated intermediate OS outcomes between Clusters 3 and 4.

**Fig 9.**
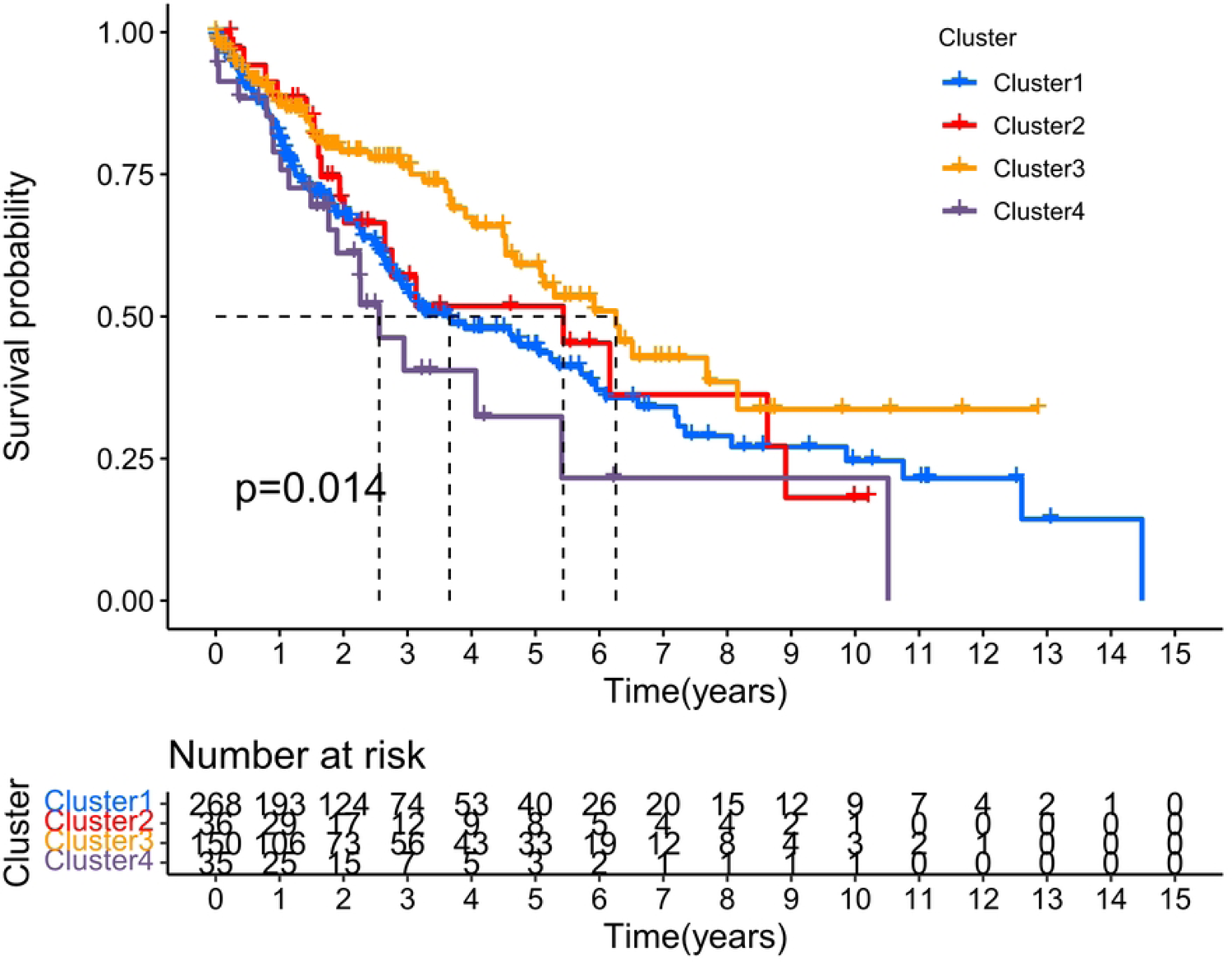
Survival analysis was performed on the four primary clusters of LUSC patients.

A heatmap was subsequently generated to illustrate the associations between EMT-related lncRNA expression, patient clinical features, and subtypes (**Fig 10**).

**Fig 10.**
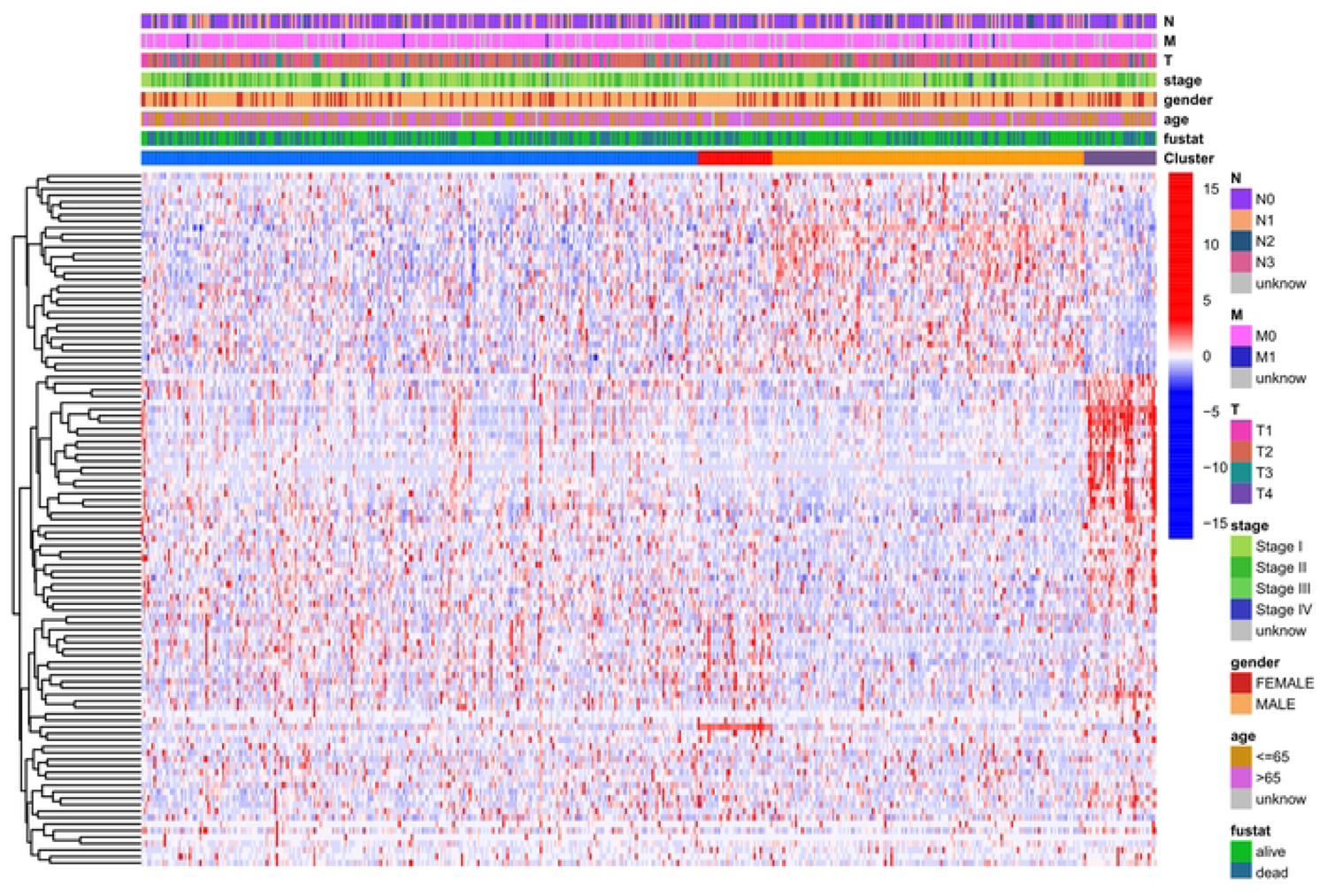
The heatmap illustrates the correlations among the four clusters, clinical characteristics, and the expression of EMT-related lncRNAs.

The clear color demarcations in the gene expression profiles of the four clusters indicated differential tumor cell behavior across the clusters. Cluster 3 predominantly consisted of T1 and T2 patients, with T3 and T4 patients being scarce. Clusters 1 and 2 displayed a more substantial increase in the proportion of T3 patients. This observation suggests that Cluster 3 may be closely related to early-stage carcinoma in situ, while Clusters 1 and 2 may be associated with tumor lesion expansion and invasion of surrounding normal tissues in LUSC. Furthermore, Cluster 4 contained a higher number of female LUSC patients.

Waterfall plots were generated using mutation data from LUSC cases in the TCGA database (**S2 Fig**). The most frequently mutated genes across all four clusters were TP53 and TTN, with mutation rates exceeding 60%. Additionally, mutation rates of CSMD in Cluster 3, CSMD and RYR2 in Cluster 4, and CSMD, SYNE1, and USH2A in Cluster 2 were all greater than 40%.

### 3.8 Immunological characteristics of distinct subtypes

Upon acquiring data on immune cell infiltration and tumor microenvironment scores for each sample, we generated a heatmap to illustrate the association between lncRNA expression, molecular subtype, and clinical factors (**Fig 11A**). The immune, stromal, and ESTIMATE scores of Clusters 2 and 3 were notably lower compared to other groups, while Cluster 4 exhibited significantly higher scores than the other three groups. In Cluster 3, the infiltration level of M0 macrophages was reduced, whereas the infiltration levels of naive B cells, follicular helper T cells, and resting dendritic cells were comparatively higher. Cluster 4 displayed elevated infiltration levels of monocytes, M2 macrophages, activated dendritic cells, resting CD4 memory T cells, neutrophils, and resting mast cells, but lower levels of M0 macrophages, M1 macrophages, eosinophils, and follicular helper T cells.

**Fig 11.**
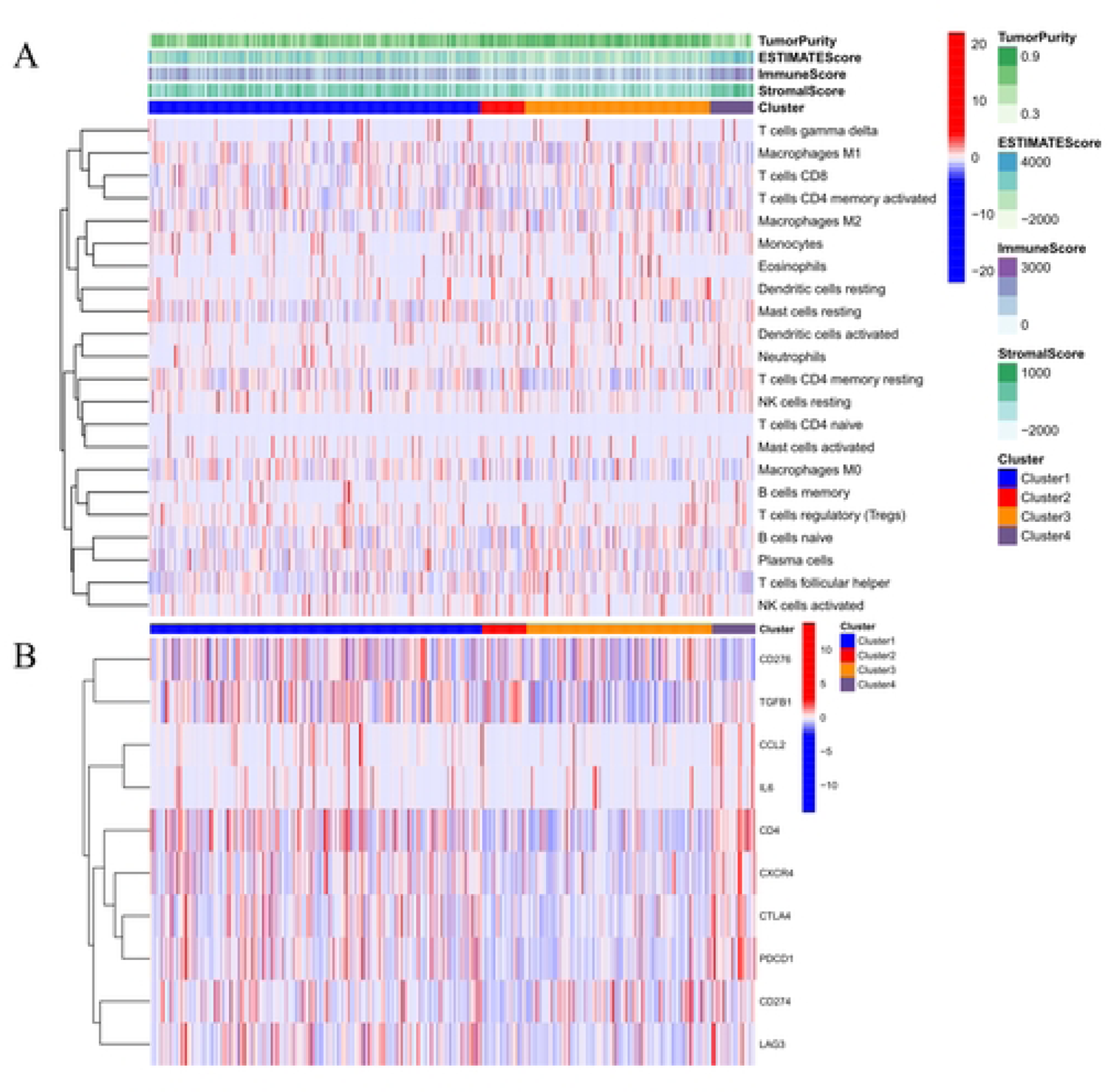
Immunological features of various clusters are depicted in the heatmaps presented in Figs A and B. Fig A illustrates the immune cell infiltration landscape and tumor microenvironment scores across four distinct clusters. Meanwhile, Fig B displays the immune checkpoint landscape across the same four clusters.

The immune checkpoint heatmap (**Fig 11B**) revealed that the highest expression levels of PDCD1, CD4, CTLA4, CXCR4, and LAG3 were observed in Cluster 4, followed by Cluster 1. Conversely, CD276 exhibited the lowest expression level in Cluster 4. The expression levels of immune checkpoint molecules in Clusters 2 and 3 were generally low. However, specifically, TGFB1 demonstrated the highest expression in Cluster 2, and CD274 displayed relatively high expression in Cluster 3.

### 3.9 Differential efficacy of immunotherapy regimens based on IPS and patient subtypes

The outcomes of the efficacy analysis, based on the Immune Prognostic Score of patients receiving distinct immunotherapy regimens, are illustrated in the violin plot below (**Fig 12**). Our findings demonstrate that the utilization of PD-1 inhibitors exhibited greater effectiveness in patients classified within Clusters 1 and 4, as opposed to other patient subtypes (**Fig 12A**). Conversely, patients categorized within Cluster 3 displayed reduced sensitivity to CTLA4 inhibitors in comparison to other subtypes (**Fig 12B**). As depicted in **Fig 12C**, the combined immunotherapy approach, incorporating both PD-1 inhibitors and CTLA4 inhibitors, proved to be more efficacious for patients in Clusters 1 and 4 relative to the administration of PD-1 inhibitors alone.

**Fig 12.**
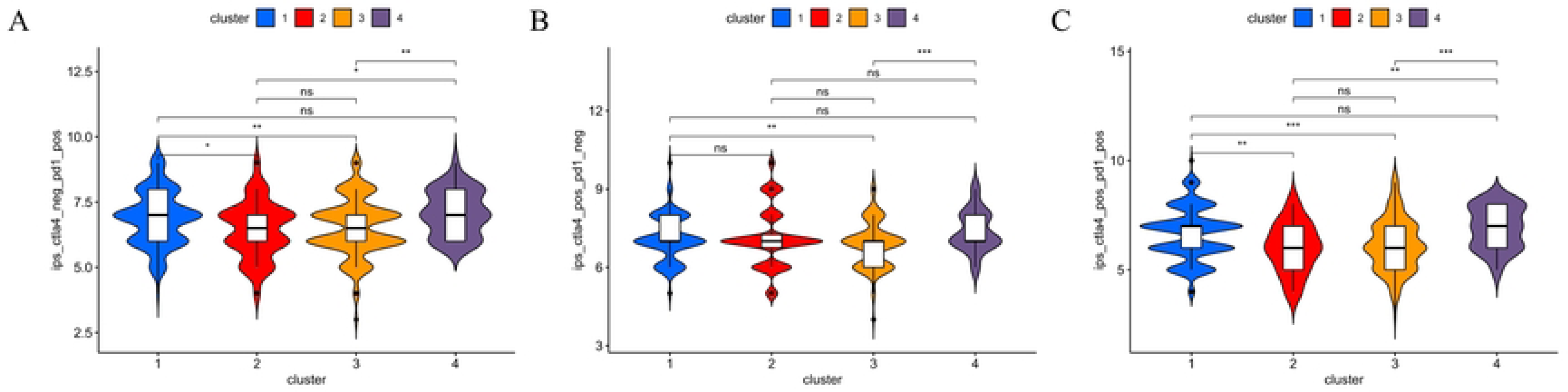
**The effectiveness of immunotherapy was evaluated in four distinct clusters**, namely PD-1 inhibitors (A), CTLA4 inhibitors (B), and the combination of PD-1 and CTLA4 inhibitors (C). The statistical significance was indicated by NS (not significant), * (P < 0.05), ** (P < 0.01), and *** (P < 0.001).

### 3.10 Prognostic implications of key EMT-LPS genes assessed through GEPIA

The Kaplan-Meier analysis revealed a significant association between increased expression levels of AC087521.1 and ST3GAL5-AS1 and decreased overall survival in patients with LUSC (*p-*value = 0.022 and *p-*value = 0.0064, respectively) (**Fig 13**). These findings suggest that AC087521.1 and ST3GAL5-AS1 may have critical roles in the pathogenesis, progression, and prognosis of LUSC. However, the expression level of AC008734.1 did not exhibit a significant correlation with OS, indicating a potential dual role of AC008734.1 in LUSC.

**Fig 13.**
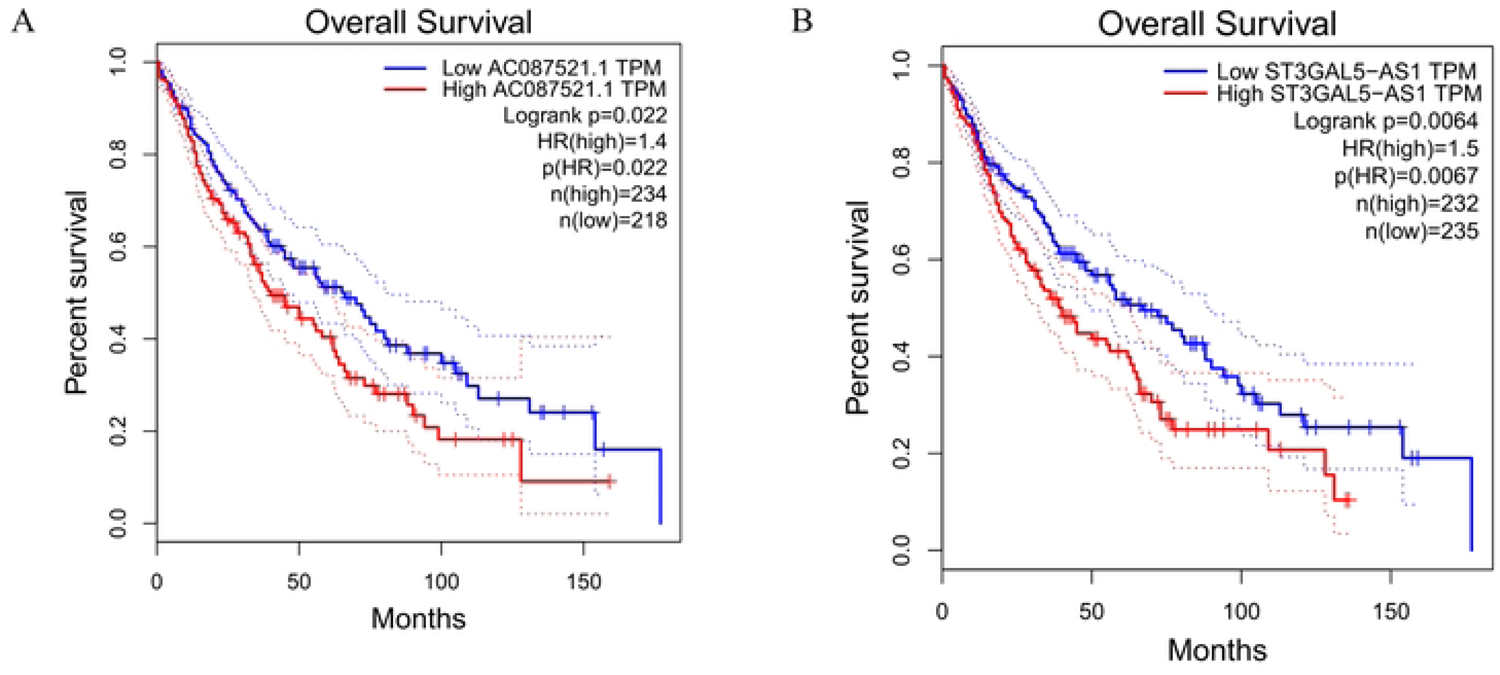
Prognostic values of key lncRNAs in the prognostic model in LUSC analyzed by GEPIA. AC087521.1 (A), ST3GAL5-AS1 (B).

## 4 Discussion

Lung squamous cell carcinoma is a major subtype of lung cancer, characterized by distinct biological pathways and prognostic factors compared to other lung cancer types (11). The exploration of novel biomarkers for prognosis prediction and the development of molecular target therapies for LUSC is of paramount importance. Emerging evidence suggests that long non-coding RNAs can promote tumor progression and metastasis by regulating epithelial-mesenchymal transition (7). However, the role of EMT-related lncRNAs in LUSC, particularly their prognostic significance, remains largely unexplored.

In this study, we constructed an EMT-related lncRNA prognostic signature based on RNA transcriptomic data and corresponding clinical information from 297 randomly selected LUSC patients in the TCGA database. We validated the EMT-LPS using a test dataset constructed from an additional 198 samples. The Kaplan-Meier survival curves and receiver operating characteristic curves of both datasets demonstrated the robust and accurate prognostic predictive ability of EMT-LPS. Independent prognostic analysis confirmed that the expression of lncRNA in EMT-LPS can serve as an independent predictor of prognosis in LUSC patients. Survival analysis across different clinical stages revealed the universal prognostic predictive value of EMT-LPS for patients with diverse clinicopathological features. We integrated patients’ pathological stage and risk score information to construct a nomogram based on these indicators, predicting 1-year, 2-year, and 3-year survival rates. The ROC curves demonstrated the robust predictive ability of the nomogram. Collectively, our study highlights the significant prognostic value of EMT-LPS and provides novel insights for precision medicine and targeted drug development for LUSC patients.

In addition, GSEA analysis revealed a plethora of significantly enriched signaling pathways in patients with higher EMT-LPS risk scores. The intricate relationship between EMT and various cellular processes, including ECM receptor interactions, focal adhesions, leukocyte transendothelial migration, and cytokine-cytokine receptor interactions, encompasses multiple signaling pathways and molecular mechanisms.

ECM receptor interactions, primarily mediated by integrins, play a pivotal role in regulating EMT Alterations in ECM composition and organization can influence EMT by modulating integrin signaling and activating downstream pathways, such as the PI3K/Akt and MAPK/ERK pathways Furthermore, EMT can impact ECM receptor expression and function, leading to changes in cell-ECM interactions and promoting cell migration and invasion (14).

Focal adhesions, specialized structures connecting the ECM to the actin cytoskeleton, are also implicated in the relationship between EMT and cell migration, invasion, and signaling (15). Key molecular players in EMT, such as the transcription factor Snail, regulate focal adhesion dynamics by modulating the expression of focal adhesion kinase (FAK) and paxillin (16). The TGF-β pathway, a potent inducer of EMT, promotes the formation of focal adhesions and stress fibers in epithelial cells (17), further emphasizing the connection between EMT and focal adhesions.

Leukocyte transendothelial migration (TEM) shares some common molecular mechanisms and signaling pathways with EMT, such as Rho GTPases and integrins, which regulate cell adhesion, migration, and cytoskeleton dynamics (18, 19). Although EMT and leukocyte TEM are involved in different physiological and pathological contexts, understanding the crosstalk between these processes may provide novel insights into cancer progression and immune response regulation (20, 21).

Cytokine-cytokine receptor interactions also play a role in the regulation of EMT. TGF-β, a key cytokine implicated in EMT, exerts its effects through binding to its receptors and activating downstream signaling pathways, such as the Smad pathway and non-Smad pathways (22, 23). Other cytokines, such as IL-6, can promote EMT through the activation of the JAK/STAT3 signaling pathway (24), while some cytokines, like IFN-γ, have been reported to inhibit EMT in certain cancer cells (25).

To recapitulate, the complex relationship between EMT and various cellular processes, including ECM receptor interactions, focal adhesions, leukocyte transendothelial migration, and cytokine-cytokine receptor interactions, involves multiple signaling pathways and molecular mechanisms. Understanding the crosstalk between these processes may provide novel therapeutic targets for the treatment of cancer.

Furthermore, the roles of several lncRNAs within our signature have been previously investigated in the context of cancer. These lncRNAs have demonstrated diverse functions in regulating tumor progression, metastasis, and response to therapy, highlighting their potential as prognostic biomarkers and therapeutic targets in oncology. AC008734.1 has been found to be closely associated with m6A and cellular apoptosis processes, and its high expression level is correlated with poor prognosis in patients with lung squamous cell carcinoma, which is consistent with our analysis results (26, 27). Ye et al. discovered that AC087521.1 is associated with poor prognosis in patients with lung adenocarcinoma (28). Li et al. used real-time quantitative PCR (RT-qPCR) to compare the expression differences of lncRNAs in human coronary artery endothelial cells (HCAECs) before and after stimulation with oxidized low-density lipoprotein (ox-LDL). They found that ST3GAL5-AS1 was significantly downregulated in HCAECs after oxLDL stimulation (29). Moreover, the mouse ortholog of ST3GAL5-AS1 was found to be differentially expressed in the kidneys of rats with immune-complex glomerulonephritis (ICGN) induced by cationic BSA (c-BSA), providing a potential lncRNA target for ICGN treatment (30). Through Chen et al.’s research, DNAH10OS is considered to be closely related to cellular autophagy and may play a crucial role in promoting breast cancer progression (31). LINC02323 is thought to promote cell growth and migration of ovarian cancer via TGF-β receptor 1 by miR-1343-3p (32). Additionally, Wu et al. found that LINC02323 facilitated the development of lung squamous cell carcinoma by miRNA sponge and RNA-binding protein dysregulation, and was linked to poor prognosis (33). lncRNA GPC5-AS1 is believed to stabilize GPC5 mRNA by competitively binding with miR-93/106a to suppress gastric cancer cell proliferation (34). Low expression of GPC5-AS1 in patients with metastatic melanoma is thought to be associated with a longer survival time (35). Enhancer RNA AC079760.2 was significantly associated with poor survival of gastric cancer patients (36). High expression of AL021368.3 is considered to be correlated with a longer survival period in patients with clear cell renal cell carcinoma (37).

In recent years, several subtype classifications of LUSC have been proposed, but a classification system with EMT as the focal point has not yet been developed. EMT plays a crucial role in tumor progression and is significantly correlated with the prognosis of LUSC. In this study, we classified 495 LUSC patients into seven subtypes based on the expression of prognosis-related EMT-related lncRNAs using the Consensus clustering algorithm. However, three of these subtypes had a small number of patients, accounting for less than 1.3% of the total studied population. To enhance the universality and scalability of our study, we excluded data from these six patients in subsequent analyses. The cumulative distribution function plot and tracking plot demonstrated the stability of our typing method.

Our innovative approach revealed that Cluster 3 patients were associated with a better prognosis and early-stage LUSC, while Cluster 4 patients generally had a worse prognosis. Given that tumor cells often hijack the EMT process to enhance tissue invasiveness and immune tolerance, we analyzed the tumor microenvironment in four patient subtypes. Cluster 4 patients exhibited a relatively high overall level of immune infiltration, potentially due to their advanced tumor progression or malignancy. Interestingly, the infiltration abundance of M2 macrophages and mast cells was also significantly higher in the tumor microenvironment of Cluster 4 patients, both of which are closely related to tumor proliferation, invasion, and migration processes (38, 39). This finding suggests that the immune system of Cluster 4 patients could exert an active antitumor effect against LUSC cells, but this effect is often suppressed by the immune escape process mediated by oncocytes.

The unique clinicopathologic features and rare driver mutations of LUSC patients present challenges in extensively benefiting from targeted therapies, such as tyrosine kinase inhibitors (TKIs) (40–42). However, the advent of immunotherapy offers a promising alternative for LUSC patients, given their complex gene mutations and high tumor mutation load (43). Immunotherapy tends to improve the prognosis of LUSC more significantly and has a lower incidence of adverse events in patients than chemotherapy alone (44). However, the clinical application of immunotherapy is currently limited, and the criteria for its application still need further refinement (5). Our study found that patients in Clusters 1 and 4 were more sensitive to immunotherapy, particularly to the combination of PD-1 inhibitors and CTLA4 inhibitors, as indicated by the IPS score in the TCIA database. This suggests that Clusters 1 and 4 patients may benefit more from combination immunotherapy, representing a novel breakthrough in the treatment of LUSC.

To synthesize the key points, our innovative classification system based on EMT-related lncRNA expression levels provides valuable insights into the prognosis and potential treatment strategies for LUSC patients. By identifying distinct patient subtypes, we can better understand the tumor microenvironment and immune infiltration patterns, ultimately paving the way for more personalized and effective immunotherapy approaches in the future.

Despite the promising results, our study has several limitations. First, the rapidly evolving field of EMT research may yield additional EMT-related genes in the future, which could further refine our findings. Second, the sample size in this study was relatively small due to the limited availability of LUSC patient data in public databases. This may have resulted in low statistical significance for some results. Additionally, the lack of detailed clinical information, such as chemotherapy regimens and drug data, precluded more in-depth analysis. Finally, the role of some EMT-related lncRNAs in non-small cell lung cancer remains unclear and warrants further investigation through in vivo or in vitro experiments.

In conclusion, this study presents an innovative EMT-associated lncRNA prognostic signature and classification system for LUSC patients, highlighting the considerable prognostic significance of EMT-LPS and the potential for personalized immunotherapy approaches based on unique patient subtypes. Despite facing certain limitations, our research lays a solid foundation for future investigations to further examine the role of EMT-related lncRNAs in LUSC and develop more effective targeted therapies to tackle this challenging cancer.

## 5 Data availability

The datasets utilized in this study can be accessed through the following repositories: TCGA (https://portal.gdc.cancer.gov/), GEPIA (http://gepia.cancer-pku.cn/), and MsigDB (https://www.gsea-msigdb.org/gsea/msigdb). These resources provide comprehensive data for the analysis conducted in the current investigation.

## 6 Supporting information

**S1 Table.** A compilation of 200 EMT-associated genes derived from the MsigDB v7.4.

**S2 Table.** A collection of 2036 EMT-associated lncRNAs acquired through correlation analysis.

**S1 Fig.** A tracking plot generated via consensus clustering algorithm analysis, examining potential EMT-related LUSC subtypes.

**S2 Fig.** Waterfall plots illustrating the genetic mutation status across the four identified clusters.

**S1 File.** The programming code employed for the development of the risk score model and the construction of molecular subtypes discussed in this investigation.

## 7 Acknowledgments

We extend our appreciation to the TCGA, GEPIA, and MsigDB databases for granting access to the essential raw research data, which has been crucial in the execution of this study.

Additionally, we would like to convey our gratitude to Dr. Yuqi Song from the Department of Thoracic Surgery, who has conducted extensive research at the University of Alberta, Canada. His invaluable assistance and guidance have substantially enhanced the coherence and scholarly rigor of our article.

## 8 Author contributions statement

XH, ZJM, and LJM contributed significantly to the conception and design of the study. GY was responsible for gathering clinical information and gene expression data. ZJM, ZBH, ZXY, and LY conducted data analysis and manuscript composition. XH, ZJM, LJM, and ZJ participated in the manuscript revision. All authors made substantial contributions to the article and approved the final submitted version.

